# Genetic profiling of *Mycobacterium tuberculosis* “modern” Beijing strains from Southwest Colombia

**DOI:** 10.1101/819425

**Authors:** Luisa Maria Nieto Ramirez, Beatriz E. Ferro, Gustavo Diaz, Richard M. Anthony, Jessica de Beer, Dick van Soolingen

## Abstract

Beijing strains of *Mycobacterium tuberculosis* (lineage2) have been associated with drug-resistance and transmission of tuberculosis worldwide. Most of the Beijing strains identified in the Colombian pacific coast exhibited a multidrug resistant (MDR) phenotype, in contrast with the phenotype observed in Beijing isolates from other South-American countries. We wanted to evaluate the clonality and genetic background of the Beijing strains isolated in Colombia that belong mostly to the spoligo-international type (SIT) 190. Out of 37 Beijing stains characterized in an 8-years period, we identified five Beijing clones; 36 that belong to the SIT190 type and only one to SIT1. Two loci in VNTR typing: MIRU 39 and QUB11b exhibited the highest level of variation among these strains. Out of the 37 Beijing strains, only one was drug susceptible, 28 represented MDR-TB, four extensively drug resistant TB (XDR-TB) and four pre-XDR-TB. The mutations *rpoB* S531L and *katG* S315T1 were the most common among the MDR strains as reported elsewhere. Whole genome sequencing analysis allowed us to classify them as modern lineage Beijing strains, sharing up to 76 out of the 275 SNPs described in Beijing strains, as identified worldwide by Schürch *et al;* including 54 non-synonymous SNPs and 23 silent mutations. We were also able to confirm the presence of 8 specific SNPs that were so far only found in the Beijing strains from Colombia. The presence of modern Beijing strains, most of them representing MDR-TB, suggests a different origin of this *M. tuberculosis* lineage compared to other Beijing strains found in neighboring countries, such as Peru. The specific 8-SNP signature confirmed the identity of these Colombian Beijing MDR strains. This work may serve as a genetic baseline to study the evolution and spread of *M. tuberculosis* Beijing strains in Colombia, which play an important role in the control of MDR-TB.

## Introduction

The Beijing genotype is considered one of the most successful and virulent lineages of *Mycobacterium tuberculosis* (*Mtb*) (1-4). Tuberculosis (TB) is still the leading cause of death due to a single infectious agent worldwide, largely because of the contribution of drug resistant *Mtb* strains (5). In many settings, Beijing strains have not only been found to be significantly correlated with drug resistance, but also with active transmission of multidrug (MDR) and extensively drug resistant (XDR) TB (1, 2, 6). Beijing is the major representative strain of lineage 2, also known as East-Asian lineage, which is defined by the deletion of region of difference (RD) RD105 (7). Two different sublineages of Beijing strains, the ancient and modern strains, have been described (also known as ‘atypical’ and ‘typical’ Beijing variants, respectively). The two variants show differences in distribution, drug resistance and virulence patterns. The ancient sub-lineage strains have been found predominantly in Russia, Korea and Japan (8-10), while modern sub-lineages are distributed worldwide, and have been largely associated with drug resistance and hypervirulence (11-13).

Although the prevalence of Beijing strains was not high in South America (14), several countries have now reported the presence of this genotype among clinical isolates (9, 15-18). The proportion of Beijing strains in Peru (close to 9%) has increased in the last decade, with predominance of the modern sub-lineage (9); at the moment without association with drug resistance. Similarly, countries such as Paraguay and Ecuador have reported Beijing strains only among pan-drug-susceptible cases.

We identified a cluster of 24 Beijing strains among MDR and XDR *Mtb* isolates from Southwestern Colombia (19, 20). In a study conducted in Buenaventura-a port city that is considered a MDR-TB hot-spot in Colombia-we found a high proportion of Beijing strains (30%) among MDR-TB isolates, we also found that 25% (18/73) of the MDR-TB cases were from previously untreated patients, suggesting active transmission of MDR-*Mtb* strains of the Beijing genotype (21) in this region. According to the SpolDB4 database, the Beijing strains observed in the Colombian pacific coast (that belong to the Spoligo-International-Type SIT 190), have been observed previously in the United States, Japan, Cuba, among other countries (22). However, more discriminative molecular analysis is needed to better characterize the cluster detected in Colombia to decipher possible epidemiological links between these cases and the worldwide spread of Beijing strains. Therefore, this study aimed to establish a comprehensive genetic profile of Beijing strains isolated in Colombia and to provide additional insights into its biology in comparison to Beijing strains from other geographical areas.

## Materials and Methods

### Ethics Statement

This study was approved by the Institutional Review Board of CIDEIM, which authorized a waiver of informed consent from the human research subjects who provided the samples. Additionally, data was anonymously analyzed and confidentiality preserved using codes instead of identifiable variables.

### Study Samples

During the period 2002 to 2010, 651 *Mtb* isolates from Valle del Cauca, Colombia were sent to CIDEIM from public and private health institutions to perform routine drug susceptibility testing (DST) on basis of the proportion Middlebrook 7H10 agar method for both first (isoniazid, rifampicin, ethambutol) and second line antibiotics (amikacin, ciprofloxacin or moxifloxacin) (23). The strains were stored at −80°C. For this study 311 were thawed to perform genetic typing and complementary genetic analyses. In total, 37 Beijing strains were detected and used for further analysis, only one isolate per patient was included.

### Genetic typing

We performed a molecular characterization of the 37 *Mtb* Beijing isolates using two Polymerase Chain Reaction (PCR) based methodologies: spoligotyping (24) and MIRU-VNTR (Mycobacteria Interspersed Repetitive Unit – Variable Number Tandem Repeat) 24 loci typing (25). DNA was extracted using the CTAB method (26). Agarose gel electrophoresis was performed to determine the number of repeats for each locus. Finally, allelic assignation was done using Quantity one software (Biorad®) to determine the length of the PCR products for each of the 24 loci analyzed. External quality control was assessed through the participation in the second worldwide proficiency study of MIRU-VNTR (27).

#### Detection of mutations associated with drug resistance

DNA was extracted according to the manufacturer’s instructions (HainLifescience GmbH, Nehren, Germany). Mutations associated with resistance to first- and second-line anti-tuberculosis drugs were detected using GenoType® MTBDR*plus* and *sl* assays (HainLifescience, Nehren, GmbH, Germany) respectively.

#### Beijing strains classification

In order to confirm that the strains belonged to the Beijing genotype family, further analyses were performed using 10 representative strains of each branch of the dendrogram based on the spoligotyping and MIRU-VNTR 24 loci typing analysis. These analyses included the evaluation of the genomic region of difference 105 (RD 105), which presence is a genetic marker for the Beijing genotype. Additionally, we performed a SNP analysis of the *fbpB*-238 region, to classify the selected Beijing strains into the Vietnam (V) “typical” (+) or atypical (–) sub-lineage, according to the scheme of Schürch *et al* (28), using a Multiplex Ligation-dependent Probe Amplification (MLPA) assay with readout facilitated by the Luminex bead technology, as described previously (29). The MLPA analysis was conducted by the Royal Tropical Institute (KIT) in The Netherlands and was also used to confirm the MIRU and the mutation analysis in the *katG, inhA, rpoB, gyrA, rrs* and *embB* resistance genes (30).

Two out of 37 Beijing strains were selected to be whole genome sequenced using Illumina sequence technology at The Broad Institute of MIT & Harvard. Analysis of SRA files was done using CLC genomics workbench 12 software, having H37Rv as the reference sequence.

### Data Analysis

The Spoligo-International-Type (SIT) number and the multiple locus VNTR analysis (MLVA) using 15 of the most discriminatory loci (MLVA MtbC15-9) were determined using the http://www.miru-vntrplus.org web page and SITVIT2 (database of the Pasteur Institute of Guadeloupe). The information obtained was used to build the phylogenetic analysis based on MIRU-VNTR patterns. VNTR typing results were recorded as character data type and compared to database at the National Institute for Public Health and the Environment (RIVM) in the Netherlands.

## Results

The identity of all 37 Beijing isolates was confirmed using spoligotyping. According to the DST, 30 of these strains were MDR-TB, 6 XDR-TB, and only one was found drug-susceptible. Patient’s median age was 29 years (IQR: 24-40 years). Most strains were isolated from female (23/37, 62%) and pulmonary TB patients (34/35, 97.1%).

All except one strain were typed as Spoligo-International number-SIT 190 (000000000003731). The remaining, susceptible strain, was typed as SIT1 (000000000003771). MIRU-VNTR analysis using 24 loci was used to explore further differences among the highly homogeneous SIT 190 cluster. However, only two loci (QUB11b and MIRU 39) were able to discriminate isolates in the SIT 190 cluster. Among those two loci, MIRU 39 revealed the highest differences, with 2, 3 and 4 repeats among the Beijing SIT190 strains. In addition, five more loci (Mtub39, Mtub04, MIRU 31, 27, and 40) showed different number of repeats between SIT 190 and SIT 1 strains (Fig 1). A phylogenetic analysis using both the spoligotyping and MIRU-VNTR 24 loci analyses confirmed the classification of SIT190 and SIT1 as different clades and showed that among the SIT190 clade there were four different clones (Fig 1).

**Fig 1.**
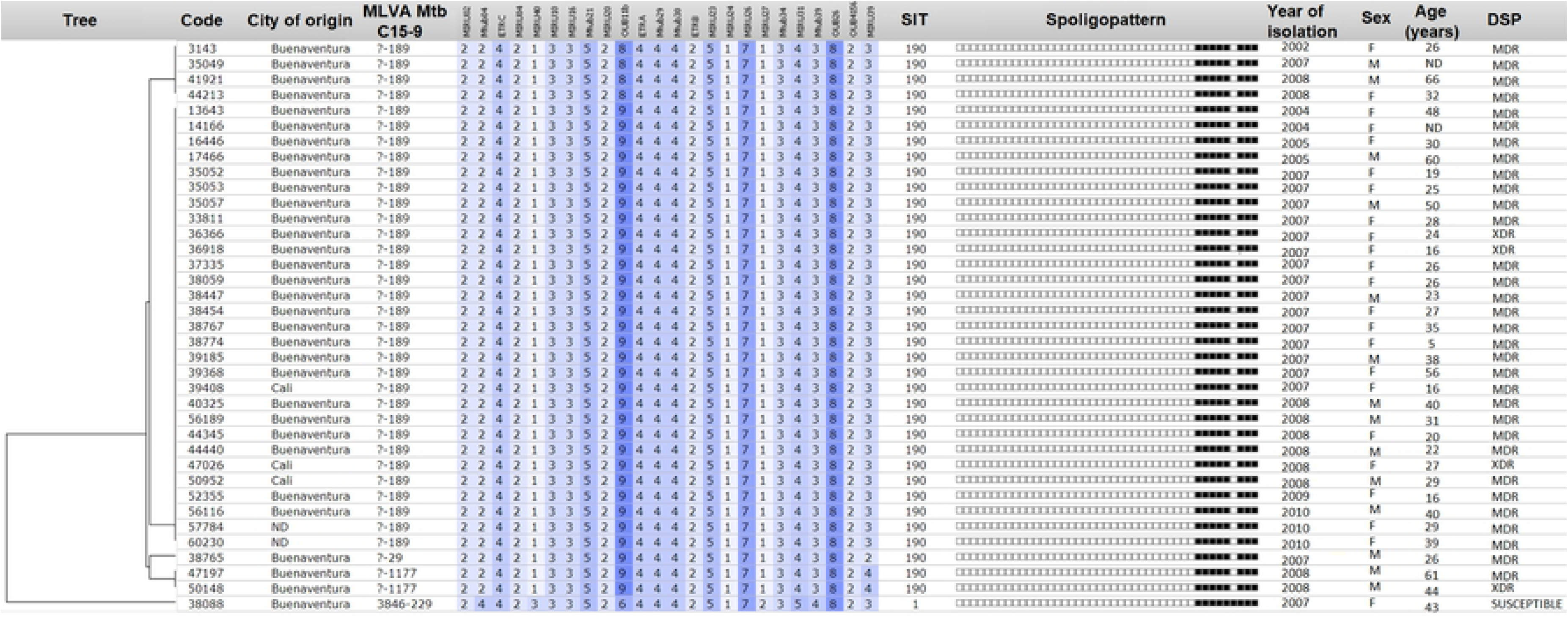
Dendrogram of the Beijing strains discriminated by MIRU VNTR 24 loci and Spoligotyping patterns. MLVA MtbC15 type for the 15 most discriminatory loci a MLVA MtbC9 type for the 9 auxiliary loci defined by Supply *et al*, 2006 (25). ND: Information not available. DSP: Drug susceptibility profile.

### Detection of mutations associated with drug resistance

In total 33 (29 MDR and 4 XDR) out of the 37 Beijing strains were successfully evaluated for the presence of mutations associated with first- and second-line drug resistance. The mutations S531L in *rpoB* and S315T in *katG*, associated with resistance to rifampicin and isoniazid, respectively, were the most frequently found as expected in MDR cases (Table 1). All four XDR strains were confirmed using the genotype analysis that also allowed the detection of four additional pre-XDR strains (defined as MDR strain that is also resistant to either a fluoroquinolone or a second-line injectable) (31). The ethambutol associated mutation M306V in the *embB* gene was also the most frequently encountered among the ethambutol resistant strains (Table 1).

**Table 1.** Frequency of mutations associated with drug resistance in Beijing genotype family strains from Colombia.

| Gene mutation probe (mutation) | % of strains (# of mutated strains/# of resistant strains detected by GenoTypeMTBDR<sub>plus</sub> and MTBDR<sub>sl</sub>) |
| --- | --- |
| *rpoB*MUT3 (S531L) | 97 (32/33) |
| *katG*MUT1 (S315T1) | 100 (33/33) |
| *inhA*MUT1 (C15T) | 3 (1/33) |
| *gyrA*MUT3C (D94G) | 13 (4/30)<sup>a</sup> |
| *rrs*MUT1 (A1401G) | 29 (8/28) |
| *embB*MUT1B (M306V) | 91 (29/32) |

<sup>a</sup> One strain had double mutations MUT3A and MUT3C.

### Beijing strains classification

The East/Asian RD105 deletion was present in all the subset of 10 Beijing strains, confirming their identification as members of the Beijing lineage. Moreover, all the strains tested, including the SIT 1 and SIT 190, were further classified into the Beijing sublineage V+.

### Whole genome sequencing analysis

The whole genome sequences of two Beijing strains from this study, designated as 38088 (fully susceptible) and 38765 (MDR), are available at the National Center for Bioinformatic Information (NCBI) with the SRA numbers SRS565195 and SRS565201 respectively. Based on the 275 SNPs proposed by Schürch *et al*., (28), these two Beijing strains isolated in Southwestern Colombia exhibited features of a modern lineage, such as the mutation at the gene *mutT2* Gly58Arg (transversion C/G at position 1286766), confirming our previous Beijing strain classification. Strain 38088 shared 75 SNPs, while the strain 38765 shared 76 SNPs out of the 275 SNPs described by Schürch *et al*. in Beijing strains identified worldwide (Table 2). Additional comparison between Beijing strains previously analyzed from Colombia and strains from this study revealed the presence of 8 additional SNPs among Beijing MDR strains (Table 3). Also, 38088 and 38765 strains have an insertion of a G at the position 1406760^1406761 in the *Rv1258c* gene.

**Table 2.** Genes (location) with single nucleotide polymorphism shared between Colombian Beijing strains and other Beijing strains characterized worldwide.

| SNP Location |  |
| --- | --- |
| Non-synonymous (n=54) | Silent (n=23) |
| *recF, fadD34, oxyS, ccsA, recD, echA4, rpsL\*, far, mutT2, narJ, fadD6, alkA, uvrC (1594906), uvrC (1595342), ansA, polA (1831220), polA (1831226), lysS, moeX, secA2, glcB, katG\*\*, rpmG, pknJ, pepE, uspA, accD1, recX, fdhD, ligB, nudC, maf, sseA, esxU, rmlC, otsA, mce4C, fadD19, ltp3, nth, ponA2, dnaQ (4156099), dnaQ (4156503), dnaQ (4155748), recR, ligC, dppB, ponA2, recR, cut5b, ligC, fadE36, embA, papA2, glpQ1, mycP2, 4393590.* | *acs, cobG, cobQ2, ctpG, cysA3\*\*\*, dnaZX, echA21, embB, fadB4, fba, fusA2, hemB, ligC, lprK, murC, ogt, polA, pyrG, radA, recC, sigM, tatD, tpiA.* |

\* Lys43Arg, found in the MDR (38765) strain only. \*\* Ser315Thr found in the MDR (38765) strain only. \*\*\*Found in the fully susceptible (38088) strain only.

**Table 3.** Comparison of the previously reported SNPs in Beijing-like strains from Colombia and the MDR Beijing strains of this study

| Accession number | Gene name | Position | Reference H37Rv | Mutation found in Beijing strains from Colombia (32) | Mutation found in MDR Beijing strain 38765 | Mutation found in FS Beijing strain 38088 |
| --- | --- | --- | --- | --- | --- | --- |
| **Rv0355c** | *PPE8* | 434226 | A | Deletion | Wild type | Wild type |
| **Rv0355c** | *PPE8* | 434227 | C | Wild type | Deletion | Wild type |
| **Rv0197** | *Rv0197* | 233949 | C | T | T | Wild type |
| **Rv0753c** | *mmsA* | 845542 | G | A | A | Wild type |
| **Rv0988** | *Rv0988* | 1106176 | A | C | C | Wild type |
| **Rv1723** | *Rv1723* | 1950068 | C | T | T | Wild type |
| **Rv2308** | *Rv2308* | 2580877 | G | T | T | T |
| **Rv2940c** | *Rv2940c* | 3279637 | G | A | A | Wild type |
| **Rv3806c** | *ubiA* | 4269304 | A | C | C | Wild type |
| **Rv3862c** | *whiB6* | 4338365 | A | G/C | G | Wild type |

FS: fully susceptible

## Discussion

In this study, we established the genetic profile of a cluster of 37 Beijing strains isolated in Colombia and compared it with Beijing strains circulating elsewhere in the world. Five interesting findings are reported: first, the conserved genetic make-up of Beijing strains isolated in Colombia confirmed our previous observation of the active and recent transmission of Beijing strains in Colombia, particularly in Buenaventura, where the main port on the Colombian pacific coast is located (19). Second, the large majority of the Beijing strains (36/37) contained mutations associated with MDR- or XDR-TB profiles, again confirming the association between Beijing and drug resistance in Colombia. Third, these Beijing strains formed a genetically highly conserved cluster, where four branches were separated, mainly based on the discriminatory capacity of 24-loci MIRU-VNTR typing (particularly two loci: 39 and QUB11b). Fourth, there was limited diversity in mutations associated with resistance in these Beijing strains, almost all were MDR due to the presence of mutations most commonly described worldwide (33-36). Finally, all the Beijing strains from this study belonged to the ‘typical’ or ‘modern’ sub-lineage, highly similar to the V+ lineages earlier described by Schürch *et al* (28), and commonly found in Vietnam. Our results support the “distribution” profile previously determined for modern Beijing strains worldwide.

Previous studies from Peru also revealed active transmission of Beijing genotype strains and the predominance of the modern sub-lineage in this country (9, 37). It has been hypothesized that Beijing strains identified in Peru probably arrived from China, when a significant Chinese migration took place, back in the 19th century (9). However, the Beijing strains circulating in Peru are genotypically different compared to the ones circulating in Colombia, indicating a different origin. The Beijing strains (mostly, SIT190) isolated in Colombia have shown highly resistant phenotypes and this varies from the Beijing strains isolated in Peru and other South American countries, where no association with drug resistance has been established (9, 14). These findings support the idea that different Beijing strains (or their ancestors) were introduced at different occasions in Southern American countries, most likely reflecting the diversity in human migration since ancient times (12). This is also in agreement with the findings of Schürch *et al* on the introduction of multiple sources of spread of Beijing strains to different geographical areas on many different occasions (28). However, it is important to highlight that the Colombian Beijing strains are genetically highly conserved and that this may reflect recent introduction and spread.

Regarding the WGS analysis, there were additional SNPs that have been also reported in other globally identified Beijing strains (28). For instance, both Beijing strains (38088 and 38765) exhibited an insertion of a G at the position 1406760^1406761 in the *Rv1258c* gene, previously identified in modern Beijing strains from Guatemala (38). Rv1258c encodes for an efflux pump that was previously proposed to be linked to streptomycin resistance (39). SNPs in *katG, rpsL* and *cysA* that were associated with antibiotic resistance, were found in the 38765 strain only. In addition, there were three non-synonymous SNPs in the genes *pta, eis* and *lipU* present in 38765. On the other hand, the synonymous SNP at the *cysA3* genes was found in 38088 strain only. Exactly the same synonymous SNP at *cysA3* was found in non-clonally distributed Beijing strains analyzed in a previous study (28) and in a modern Beijing strains from Guatemala (38).

Except the frameshift deletion at the position 434226, all the remaining SNPs previously reported as exclusively present in the Beijing-like strains from Colombia (32) were also found in the MDR-TB Beijing strain 38765 that was subjected to WGS in this study (Table 2). Instead of the deletion at the position 434226, our Beijing MDR-TB strain (38765) also had a deletion just in the next nucleotide position (434227) (Table 3). For the FS Beijing strain (38088), only one mutation at the *Rv2308* gene was shared with another Beijing-like strains from Colombia as well as with 38765 (Table 3). *Rv2308* encodes for a conserved hypothetical protein that may act as a transcriptional regulator (40). Importantly, this SNP found in Rv2308 has not previously identified in other Beijing strains worldwide (28).

We acknowledge several limitations in the present study. The study design included convenience sampling with a limited number of Beijing isolates that were available in a repository in Colombia. Nevertheless, the samples included all Beijing strains found in that setting during a 10 years period. Additionally, the repository was built over the years with the cultures successfully sent to our institution for surveillance purposes.

Another potential limitation of our study is related to the selection of SNPs to characterize a population of Beijing strains which is still controversial; there is no consensus on which SNPs are most informative and reliable to accurately describe the phylogeny of this important genotype family of *M. tuberculosis*. For that reason, here we utilized the SNPs used in previous work (28), as this represented one of the most extensive studies in this field. However, there is still a need to harmonize the SNPs as well as the set of MIRU loci to subdivide Beijing strains, in order to strengthen the data analysis and its impact. Information obtained by high throughput technologies, like whole genome sequencing, will shortly improve our understanding and definition of locally and worldwide circulating Beijing strains, but will not be readily available in all settings.

In conclusion, Beijing strains that circulated in Colombia from 2002 to 2010 belong to a genetically conserved cluster of modern Beijing strains, and have important phenotypic differences (being most of them MDR strains) from the strains circulating in neighboring countries like Peru. Continuing with the surveillance and monitoring of the local *Mtb* structure remains important in understanding the epidemiology of MDR-TB and facilitate the development of control strategies against (MDR) TB.

## Acknowledgements

This work was supported by local health authorities from Valle del Cauca State and the city of Santiago de Cali. We would also want to thank Juan Carlos Rozo from CIDEIM, Sarah Sangstake and Indra Bergval from KIT Amsterdam for their technical support. Additionally, we want to thank Dr. Ashlee Earl and the Broad Institute for the whole genome sequencing analysis.

## Author contributions

LMN*: Designed and performed experiments, analyzed data and wrote drafts.

BEF*: Designed and performed experiments, analyzed data and wrote drafts.

GD: Conducted laboratory analysis and analyzed data,

JdB: Conducted laboratory analysis, reviewed and edited drafts.

RA: Designed and performed experiments, analyzed data, reviewed and edited drafts.

DvS: Designed and performed experiments, analyzed data, reviewed and edited drafts.

## References

1. Glynn JR, Whiteley J, Bifani PJ, Kremer K, van Soolingen D. Worldwide occurrence of Beijing/W strains of Mycobacterium tuberculosis: a systematic review. Emerg Infect Dis. 2002;8(8):843–9.

2. Parwati I, van Crevel R, van Soolingen D. Possible underlying mechanisms for successful emergence of the Mycobacterium tuberculosis Beijing genotype strains. Lancet Infect Dis. 2010;10(2):103–11.

3. Buu TN, Huyen MN, Lan NT, Quy HT, Hen NV, Zignol M, et al. The Beijing genotype is associated with young age and multidrug-resistant tuberculosis in rural Vietnam. Int J Tuberc Lung Dis. 2009;13(7):900–6.

4. European Concerted Action on New Generation Genetic Markers and Techniques for the Epidemiology and Control of Tuberculosis. Beijing/W genotype Mycobacterium tuberculosis and drug resistance. Emerg Infect Dis. 2006;12(5):736–43.

5. World, Health, Organization. Global Tuberculosis Report 2017.

6. Bifani PJ, Mathema B, Kurepina NE, Kreiswirth BN. Global dissemination of the Mycobacterium tuberculosis W-Beijing family strains. Trends Microbiol. 2002;10(1):45–52.

7. Shitikov E, Kolchenko S, Mokrousov I, Bespyatykh J, Ischenko D, Ilina E, et al. Evolutionary pathway analysis and unified classification of East Asian lineage of Mycobacterium tuberculosis. Sci Rep. 2017;7(1):9227.

8. Mokrousov I, Ly HM, Otten T, Lan NN, Vyshnevskyi B, Hoffner S, et al. Origin and primary dispersal of the Mycobacterium tuberculosis Beijing genotype: clues from human phylogeography. Genome Res. 2005;15(10):1357–64.

9. Iwamoto T, Grandjean L, Arikawa K, Nakanishi N, Caviedes L, Coronel J, et al. Genetic diversity and transmission characteristics of Beijing family strains of Mycobacterium tuberculosis in Peru. PLoS One. 2012;7(11):e49651.

10. Liu Q, Luo T, Dong X, Sun G, Liu Z, Gan M, et al. Genetic features of Mycobacterium tuberculosis modern Beijing sublineage. Emerg Microbes Infect. 2016;5:e14.

11. de Keijzer J, de Haas PE, de Ru AH, van Veelen PA, van Soolingen D. Disclosure of selective advantages in the “modern” sublineage of the Mycobacterium tuberculosis Beijing genotype family by quantitative proteomics. Mol Cell Proteomics. 2014;13(10):2632–45.

12. Comas I, Coscolla M, Luo T, Borrell S, Holt KE, Kato-Maeda M, et al. Out-of-Africa migration and Neolithic coexpansion of Mycobacterium tuberculosis with modern humans. Nat Genet. 2013;45(10):1176–82.

13. Ribeiro SC, Gomes LL, Amaral EP, Andrade MR, Almeida FM, Rezende AL, et al. Mycobacterium tuberculosis strains of the modern sublineage of the Beijing family are more likely to display increased virulence than strains of the ancient sublineage. J Clin Microbiol. 2014;52(7):2615–24.

14. Ritacco V, López B, Cafrune PI, Ferrazoli L, Suffys PN, Candia N, et al. Mycobacterium tuberculosis strains of the Beijing genotype are rarely observed in tuberculosis patients in South America. Mem Inst Oswaldo Cruz. 2008;103(5):489–92.

15. Laserson KF, Osorio L, Sheppard JD, Hernández H, Benitez AM, Brim S, et al. Clinical and programmatic mismanagement rather than community outbreak as the cause of chronic, drug-resistant tuberculosis in Buenaventura, Colombia, 1998. Int J Tuberc Lung Dis. 2000;4(7):673–83.

16. Abadía E, Sequera M, Ortega D, Méndez MV, Escalona A, Da Mata O, et al. Mycobacterium tuberculosis ecology in Venezuela: epidemiologic correlates of common spoligotypes and a large clonal cluster defined by MIRU-VNTR-24. BMC Infect Dis. 2009;9:122.

17. Candia N, Lopez B, Zozio T, Carrivale M, Diaz C, Russomando G, et al. First insight into Mycobacterium tuberculosis genetic diversity in Paraguay. BMC Microbiol. 2007;7:75.

18. Jiménez P, Calvopiña K, Herrera D, Rojas C, Pérez-Lago L, Grijalva M, et al. [Identification of the Mycobacterium tuberculosis Beijing lineage in Ecuador]. Biomedica. 2017;37(2):233–7.

19. Ferro BE, Nieto LM, Rozo JC, Forero L, van Soolingen D. Multidrug-resistant Mycobacterium tuberculosis, Southwestern Colombia. Emerg Infect Dis. 2011;17(7):1259–62.

20. Nieto LM, Ferro BE, Villegas SL, Mehaffy C, Forero L, Moreira C, et al. Characterization of extensively drug-resistant tuberculosis cases from Valle del Cauca, Colombia. J Clin Microbiol. 2012;50(12):4185–7.

21. Villegas SL, Ferro BE, Perez-Velez CM, Moreira CA, Forero L, Martínez E, et al. High initial multidrug-resistant tuberculosis rate in Buenaventura, Colombia: a public-private initiative. Eur Respir J. 2012;40(6):1569–72.

22. Brudey K, Driscoll JR, Rigouts L, Prodinger WM, Gori A, Al-Hajoj SA, et al. Mycobacterium tuberculosis complex genetic diversity: mining the fourth international spoligotyping database (SpolDB4) for classification, population genetics and epidemiology. BMC Microbiol. 2006;6:23.

23. Kent PT, Kubica GP. Public Health Mycobacteriology. A guide for the level III laboratory. U.S Department of Health and Human Services PHS, Centers for Disease Control CDC, editor. Atlanta, GA1985.

24. Kamerbeek J, Schouls L, Kolk A, van Agterveld M, van Soolingen D, Kuijper S, et al. Simultaneous detection and strain differentiation of Mycobacterium tuberculosis for diagnosis and epidemiology. J Clin Microbiol. 1997;35(4):907–14.

25. Supply P, Allix C, Lesjean S, Cardoso-Oelemann M, Rüsch-Gerdes S, Willery E, et al. Proposal for standardization of optimized mycobacterial interspersed repetitive unit-variable-number tandem repeat typing of Mycobacterium tuberculosis. J Clin Microbiol. 2006;44(12):4498–510.

26. van Embden JD, Cave MD, Crawford JT, Dale JW, Eisenach KD, Gicquel B, et al. Strain identification of Mycobacterium tuberculosis by DNA fingerprinting: recommendations for a standardized methodology. J Clin Microbiol. 1993;31(2):406–9.

27. de Beer JL, Ködmön C, van Ingen J, Supply P, van Soolingen D. Second worldwide proficiency study on variable number of tandem repeats typing of Mycobacterium tuberculosis complex. Int J Tuberc Lung Dis. 2014;18(5):594–600.

28. Schürch AC, Kremer K, Hendriks AC, Freyee B, McEvoy CR, van Crevel R, et al. SNP/RD typing of Mycobacterium tuberculosis Beijing strains reveals local and worldwide disseminated clonal complexes. PLoS One. 2011;6(12):e28365.

29. Bergval I, Sengstake S, Brankova N, Levterova V, Abadía E, Tadumaze N, et al. Combined species identification, genotyping, and drug resistance detection of Mycobacterium tuberculosis cultures by MLPA on a bead-based array. PLoS One. 2012;7(8):e43240.

30. Sengstake S, Bablishvili N, Schuitema A, Bzekalava N, Abadia E, de Beer J, et al. Optimizing multiplex SNP-based data analysis for genotyping of Mycobacterium tuberculosis isolates. BMC Genomics. 2014;15:572.

31. Banerjee R, Allen J, Westenhouse J, Oh P, Elms W, Desmond E, et al. Extensively drug-resistant tuberculosis in california, 1993-2006. Clin Infect Dis. 2008;47(4):450–7.

32. Rodríguez-Castillo JG, Pino C, Niño LF, Rozo JC, Llerena-Polo C, Parra-López CA, et al. Comparative genomic analysis of Mycobacterium tuberculosis Beijing-like strains revealed specific genetic variations associated with virulence and drug resistance. Infect Genet Evol. 2017;54:314–23.

33. Ferro BE, García PK, Nieto LM, van Soolingen D. Predictive value of molecular drug resistance testing of Mycobacterium tuberculosis isolates in Valle del Cauca, Colombia. J Clin Microbiol. 2013;51(7):2220–4.

34. Luo T, Zhao M, Li X, Xu P, Gui X, Pickerill S, et al. Selection of mutations to detect multidrug-resistant Mycobacterium tuberculosis strains in Shanghai, China. Antimicrob Agents Chemother. 2010;54(3):1075–81.

35. Devaux I, Manissero D, Fernandez de la Hoz K, Kremer K, van Soolingen D, network E. Surveillance of extensively drug-resistant tuberculosis in Europe, 2003-2007. Euro Surveill. 2010;15(11).

36. Café Oliveira LN, Muniz-Sobrinho JaS, Viana-Magno LA, Oliveira Melo SC, Macho A, Rios-Santos F. Detection of multidrug-resistant Mycobacterium tuberculosis strains isolated in Brazil using a multimarker genetic assay for katG and rpoB genes. Braz J Infect Dis. 2016;20(2):166–72.

37. Sheen P, Couvin D, Grandjean L, Zimic M, Dominguez M, Luna G, et al. Genetic diversity of Mycobacterium tuberculosis in Peru and exploration of phylogenetic associations with drug resistance. PLoS One. 2013;8(6):e65873.

38. Saelens JW, Lau-Bonilla D, Moller A, Medina N, Guzmán B, Calderón M, et al. Whole genome sequencing identifies circulating Beijing-lineage Mycobacterium tuberculosis strains in Guatemala and an associated urban outbreak. Tuberculosis (Edinb). 2015;95(6):810–6.

39. Villellas C, Aristimuño L, Vitoria MA, Prat C, Blanco S, García de Viedma D, et al. Analysis of mutations in streptomycin-resistant strains reveals a simple and reliable genetic marker for identification of the Mycobacterium tuberculosis Beijing genotype. J Clin Microbiol. 2013;51(7):2124–30.

40. Camus JC, Pryor MJ, Médigue C, Cole ST. Re-annotation of the genome sequence of Mycobacterium tuberculosis H37Rv. Microbiology. 2002;148(Pt 10):2967–73.

